# Shared mechanisms of dopamine and ATP transmission in the nucleus accumbens

**DOI:** 10.64898/2026.03.24.713678

**Authors:** SC Linderman, LH Ford, JD Dickerson, C Ahrens, HA Wadsworth, SC Steffensen, JT Yorgason

## Abstract

Dopamine (DA) neurons of the midbrain project throughout the striatum, including the nucleus accumbens core (NAc) and are thought to co-release ATP with DA from vesicles. The mechanisms of evoked NAc ATP release and clearance and their relationship to exocytotic DA transmission are largely unexplored and the focus of the present work. Using fast scan cyclic voltammetry (FSCV), we measured simultaneous ATP and DA transmission in response to pharmacological manipulations of release and reuptake cellular machinery. ATP transmission is tightly coupled to that of DA, though ATP release concentrations are typically smaller. Manipulations that increase DA transmission (increased release via 4-aminopyridine Kv channel blockade or decreased uptake via cocaine) also increase ATP transmission, though to a smaller extent. Blocking DA vesicular packaging (reserpine) or action potentials (lidocaine), results in attenuated DA and ATP release. Interestingly, reserpine or lidocaine can result in completely abolished DA release, but not a complete prevention in ATP release, suggesting a secondary source for ATP transmission that’s not dependent on DA terminals. Both transmitters were reduced to a similar extent following nAChR blockade, demonstrating that nAChR activation regulates ATP in addition to DA. Surprisingly, cocaine inhibition of DATs reduced clearance for both ATP and DA, which correlated with one another when cocaine concentration was highest. There was also a strong relationship between the effect of cocaine on release of ATP and DA. As the first FSCV study to examine evoked NAc ATP release, this paper bridges prior work to confirm the strong association between ATP and DA in the mesolimbic circuit and identifies unexpected overlap in mechanisms regulating their transmission. Our results contribute novel evidence of both vesicular and non-vesicular ATP release in the NAc and demonstrate that extracellular ATP is a modulator of DA terminal function.

## Introduction

Adenosine triphosphate (ATP), generated via oxidative phosphorylation in cell mitochondria3/24/2026 5:55:00 AM (Bonora et al., 2012; Erecińska & Silver, 1989), is widely known for its role in muscle contraction and protein function (Bonora et al., 2012; Erecińska & Silver, 1989; Rall, 2023). In the brain, it is essential for neuronal function—the sodium-potassium pump alone is singlehandedly responsible for roughly 55% of the brain’s energy consumption (Engl & Attwell, 2015). Besides being directly essential for neuronal signaling via active ion exchange pumps, ATP is also a robust central nervous system (CNS) signaling molecule, involved in sleep, immune function, and vasodilation. Acting on purinergic receptors (e.g. A1, A2A, A2B, A3, P2X, and P2Y subtypes), ATP can convey excitatory and/or inhibitory information to nearby cells in the brain (Adamah-Biassi et al., 2015; Chang et al., 2020) depending upon cellular signaling systems. Prior research has yielded evidence of vesicular exocytosis of ATP, which further supports this neurotransmitter-like role (Borgus et al., 2021; Cisneros-Mejorado et al., 2015; Hasuzawa et al., 2020; Ho et al., 2015).

The mesolimbic dopamine (DA) system, originating in the ventral tegmental area (VTA) and projecting to the Nucleus Accumbens core (NAc), is implicated in reward/reinforcement learning (Moore & Bloom, 1978). Recent immunohistochemistry experiments have identified VTA DA neurons as having high expression of vesicular nucleotide transporter (VNUT), which packages ATP into vesicles (Ho et al., 2015). However, it is unknown how ATP signaling contributes to mesolimbic circuitry. The current literature surrounding ATP as a neurotransmitter is broad but still largely unexplored, though it is phylogenetically one of the oldest signaling molecules (Plattner & Verkhratsky, 2016). To date, several different mechanisms of cellular ATP release have been identified from a variety of neuronal cell types. In mammalian taste cells, ATP is passively diffused through membrane-bound, ATP-permeable ion channels (Romanov et al., 2008). In the retina, ATP efflux can be similarly mediated by conductive diffusion, but active exocytosis via synaptic vesicles has also been demonstrated, which requires the packaging of ATP into vesicles via the VNUT (Ho et al., 2015). In chromaffin cells, ATP enhances catecholamine packaging (Larsson et al., 2019; Majdi et al., 2019). As mentioned, VNUT expression has also been observed in midbrain VTA and substantia nigra DA neurons (Ho et al., 2015). Further, the caudate-putamen, which receives its DA inputs largely from the substantia nigra, has been reported to exhibit release of evoked and spontaneous DA release alongside ATP/adenosine (Borgus et al., 2021; Puthongkham et al., 2020). Caudate DA release is sensitive to modulation by adenosine receptor activity (Borgus et al., 2021), and we and others have observed ATP receptors throughout the NAc (Wadsworth et al., 2024). Currently, the mechanisms governing NAc ATP release and clearance remain unexamined. Considering the abundance of VNUT in VTA DA neurons as well as the expression of ATP receptors in the NAc, where they project, the present work examined ATP transmission and corresponding roles for regulators of DA release and clearance. Fast-scan cyclic voltammetry (FSCV) is a powerful technique capable of measuring subsecond ATP/adenosine release (Nguyen & Venton, 2015; Ross & Venton, 2014) while concurrently measuring DA release (Everett et al., 2022; Yorgason et al., 2011) through the use of a modified scanning voltage. Thus, FSCV was used to test the hypothesis that ATP transmission is regulated by DA related mechanisms.

## Materials and Methods

### Animal Care

Male and female transgenic mice on a C57BL/6J background were given *ad libitum* access to food and water and maintained on a 12:12 hour light/dark cycle. All protocols and animal care procedures were in accordance with the National Institutes of Health *Guide for the care and use of laboratory animals* and approved by the Brigham Young University Animal Care and Use Committee.

### Brain Slice Preparation

Mice were anesthetized with isoflurane (CHEBI:6015; Patterson Veterinary) before being decapitated with a razor blade and their brains carefully placed in pre-oxygenated (95% O2/5% CO2) artificial cerebral spinal fluid (ACSF; 35°, pH ∼7.4). The ACSF consisted of (in mM) as follows: 126 NaCl, 2.5 KCl, 1.2 NaH2PO4, 1.2 MgCl2, 21.4 NaHCO3, and 11 D-glucose. The ACSF used for slicing was treated with ketamine (2 mM), an NMDA receptor antagonist, to prevent excitotoxity while the brains were sliced with a vibratome (Leica VT1000 S) for a thickness of 220 micrometers.

### Drug Preparation and Administration

The following concentrations of drugs, obtained from Sigma-Aldrich and Tocris Bioscience, were bath applied for ∼50 minutes during FSCV experiments where specified: 4-aminopyridine (4-AP; 30 μM; Kv channel blocker), lidocaine (100 μM; NaV channel blocker), quinpirole (100 nM; D2R agonist), sulpiride (600 nM; D2R antagonist), reserpine (1 µM; VMAT inhibitor), hexamethonium (HEX; 200 μM; nAChR antagonist), and cocaine (30 μM DAT antagonist and sodium channel blocker).

### Fast-Scan Cyclic Voltammetry (FSCV)

Dopamine and ATP release were detected concurrently using previously established FSCV protocols (Borgus et al., 2021; Nguyen & Venton, 2015; Yorgason et al., 2011). Briefly, a stimulating borosilicate glass electrode (P-87 Horizontal pipette puller; Sutter Instruments) was filled (1 mM KCl) and microscopically inserted in the brain region of interest with electronic micromanipulators (Siskiyou) at ∼85 μm below the slice surface. The recording electrode was a carbon fiber electrode (CFE) made by aspirating a carbon fiber (diameter ∼7 μm, Thornel T-650, Cytec) into a borosilicate glass capillary tube (TW150, World Precision Instruments), before heating and pulling on a horizontal puller to form a tight seal of glass around the carbon fiber. The carbon fiber was then trimmed (∼100-200 μm), filled with KCl (1 mM), and placed ∼150 μm away from the stimulating electrode. A voltage ramp was applied to the CFE (−0.4 to +1.5 V) at a scan rate of 400 V/sec and a frequency of 10 Hz to measure DA and ATP corelease, with all measurements being made against a Ag/AgCl reference electrode (Borgus et al., 2021). Demon Voltammetry and Analysis software (Yorgason et al., 2011) was used to automate acquisition and facilitate analysis. Experiments measured neurotransmitter release across a period of 20-50 min to establish stable ACSF baselines. Dopamine and ATP release was evoked through electrical stimulation (1 pulse every 5 min or 2 min for HEX and cocaine experiments) with the stimulating electrode (monophasic+, 0.5 msec; 30-100 nA).

### Data Analysis and Statistical Analysis

Evoked DA and ATP release and reuptake were analyzed using DEMON Voltammetry software (Yorgason et al., 2011). Current was measured at peak oxidation for both DA and ATP (DA: 0.6 V, ATP: 1.3/1.0 V). The presence of an analyte was manually evaluated using the false color plot, current tracing, and CV graph data. Dopamine calibration has been previously described (Everett et al., 2022). ATP calibrations were performed similarly. Briefly, bath administration of known concentrations (1-10 µM) of DA or ATP were recorded, and the calibration factors for CFEs calculated and compared to the background current amplitude to establish a linear regression model for calibration values. To compare release and uptake values across multiple animals and slices, DA and ATP FSCV signals were averaged (within subject) across the last three ACSF recordings to establish a baseline value that subsequent DA and ATP signals were normalized to. Thus, post-drug conditions were compared to pre-drug baseline conditions in the same slice for a within-subject experimental design. An additional pre/post analysis (Fig. 6) was also performed for cocaine experiments using earlier recordings (while the concentration of cocaine was still increasing in the slice bath) to investigate effects of cocaine while it exhibited enhancement of DA release (Venton et al., 2006). The concentration of cocaine in this analysis is unknown but estimated to be between 1-10 µM based on transmission kinetics parameters (Everett et al., 2022). The GraphPad outlier calculator was used to perform Grubbs’ test, the detection of significant outliers in a given data set. The Shapiro-Wilk test was used to confirm normality where data appeared highly non-normal. Statistical analysis was performed using Prism 5 (GraphPad). Significance for all tests was set at *p*L<L0.05. Values are expressed as meanL±LSEM. Significance levels are indicated on graphs with asterisks *,**,***, corresponding to significance levels *p*L<L0.05, 0.01 and 0.001, respectively. Data will be shared on a per request basis.

## Results

### Dopamine and ATP Release and Clearance are Related

We set out to characterize the release and reuptake of ATP by using FSCV to detect extracellular ATP and relate to simultaneously detected DA in coronal NAc slices (**Fig. 1**). NAc brain slices were obtained and stimulated using this extended waveform. Example traces and corresponding colorplot for co-detection of evoked ATP and DA signals in NAc brain slices is shown (**Fig. 1A,B**). Calibrations were performed in a flow cell using the extended waveform to confirm ATP and DA detection (**Fig. 1C**) with DA oxidizing around 0.6 V and ATP oxidizing around 1.5 V (Borgus et al., 2021; Yorgason et al., 2011). The mean amplitude for evoked ATP and DA release was 208.1 ± 30.68 nM and 410.6 ± 52.95 nM, respectively (**Fig. 1D**; Amplitude: two-tailed paired t-test, *t* _28_=3.637, *p*=0.0011). Rate of release (maximal upward velocity) of ATP was 97 nM/sec slower than DA (**Fig. 1E**; two-tailed paired t-test, t_29_=5.033, *p<*0.0001). The clearance rate was similar between the two neurotransmitters (**Fig. 1F**; Tau: two-tailed paired t-test, *t*_29_=0.6182, *p*=0.5413). There was a significant positive correlation between the release of neurotransmitters (**Fig. 1G**; Spearman’s r=0.5995, *p*<0.0006) and the clearance of neurotransmitters (**Fig. 1H**; Spearman’s r=0.7335, *p*<0.0001). The strong relationship between ATP and DA release and clearance suggests that the mechanisms underlying these phenomena are tightly coupled.

**Figure 1:**
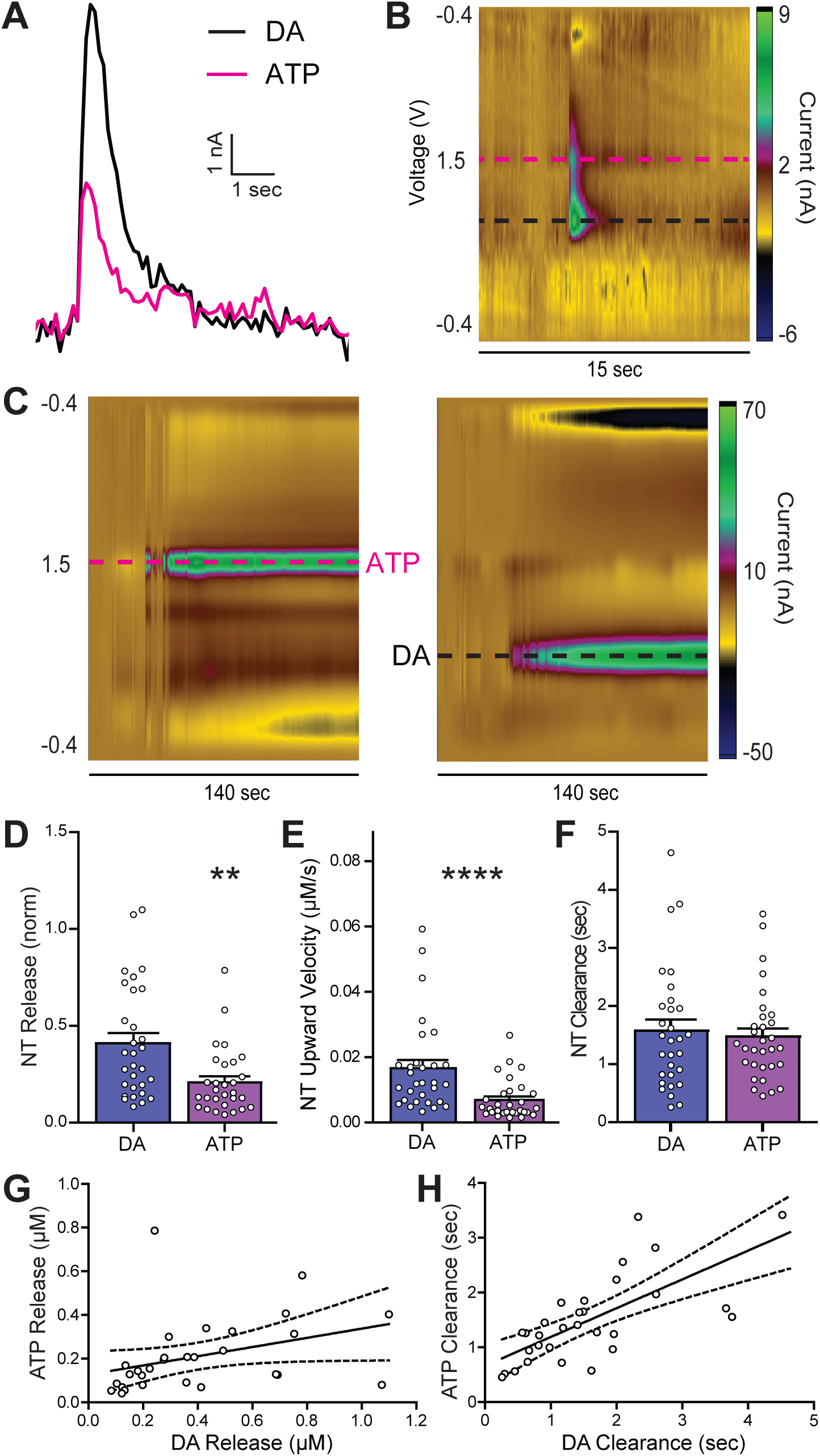
Characterization of Evoked NAc DA and ATP Release. **A** Representative traces of evoked DA (black) and ATP (pink) release. **B** Representative color plot showing DA and ATP release indicated by dashed lines (black and pink, respectively). **C** Cyclic voltammogram showing DA and ATP release indicated by arrows (black and pink, respectively). **D** Evoked DA and ATP release mean amplitudes are significantly different (two-tailed paired t-test). **E** Evoked DA and ATP have significant differences in maximal upward velocity. **F** No difference in clearance time between evoked DA and ATP (two-tailed paired t-test). **G** The correlation between ATP and DA release is significant (*p*<0.001). **H** The correlation between ATP and DA clearance rates is significant (*p*<0.001). Asterisks ** indicate significance level *p*<0.01 compared to DA Peak Height.

### DA and ATP Release are Action Potential Dependent

ATP release mechanisms in the NAc have not been characterized, but DA release is well known to exhibit dependency on voltage-gated potassium and sodium channel activity (Yorgason et al., 2017). Bath application of the voltage-gated potassium (Kv) channel blocker 4-AP (30 μM) enhanced both DA and ATP release (**Fig. 2A,B**), while subsequent application of the voltage-gated sodium (NaV) channel blocker lidocaine (100 μM) inhibited both DA and ATP release (**Fig. 2A,C**). 4-aminopyridine increased the amplitude of evoked DA and ATP release, with significant main effect of 4-AP for each neurotransmitter, and no significant difference in the response between neurotransmitters, nor interaction between these comparisons (**Fig. 2B**; two-way repeated-measures ANOVA; 4AP: F_1,10_=8.267, *p*=0.0165; neurotransmitter: F_1,10_=0.0493, *p*=0.828; interaction: F_1,10_=0.0873, *p*=0.7737). Lidocaine significantly decreased the amplitude of both DA and ATP release, with no main effect of neurotransmitter type or interaction between these variables (**Fig. 2C**; two-way ANOVA; Lidocaine: F_1,19_=34.61, *p*<0.0001; neurotransmitter: F_1,19_=2.425, *p*=0.1359; interaction: F_1,19_=1.890, *p*=0.1852; Bonferroni post hoc, *p*<0.001 for DA, *p*<0.01 for ATP). Notably, some ATP signals were not sensitive to lidocaine, suggesting that ATP release can also occur through a non-action potential mediated pathway. However, for the most part, ATP and DA release appear to share common action-potential related mechanisms in Kv and NaV channels.

**Figure 2:**
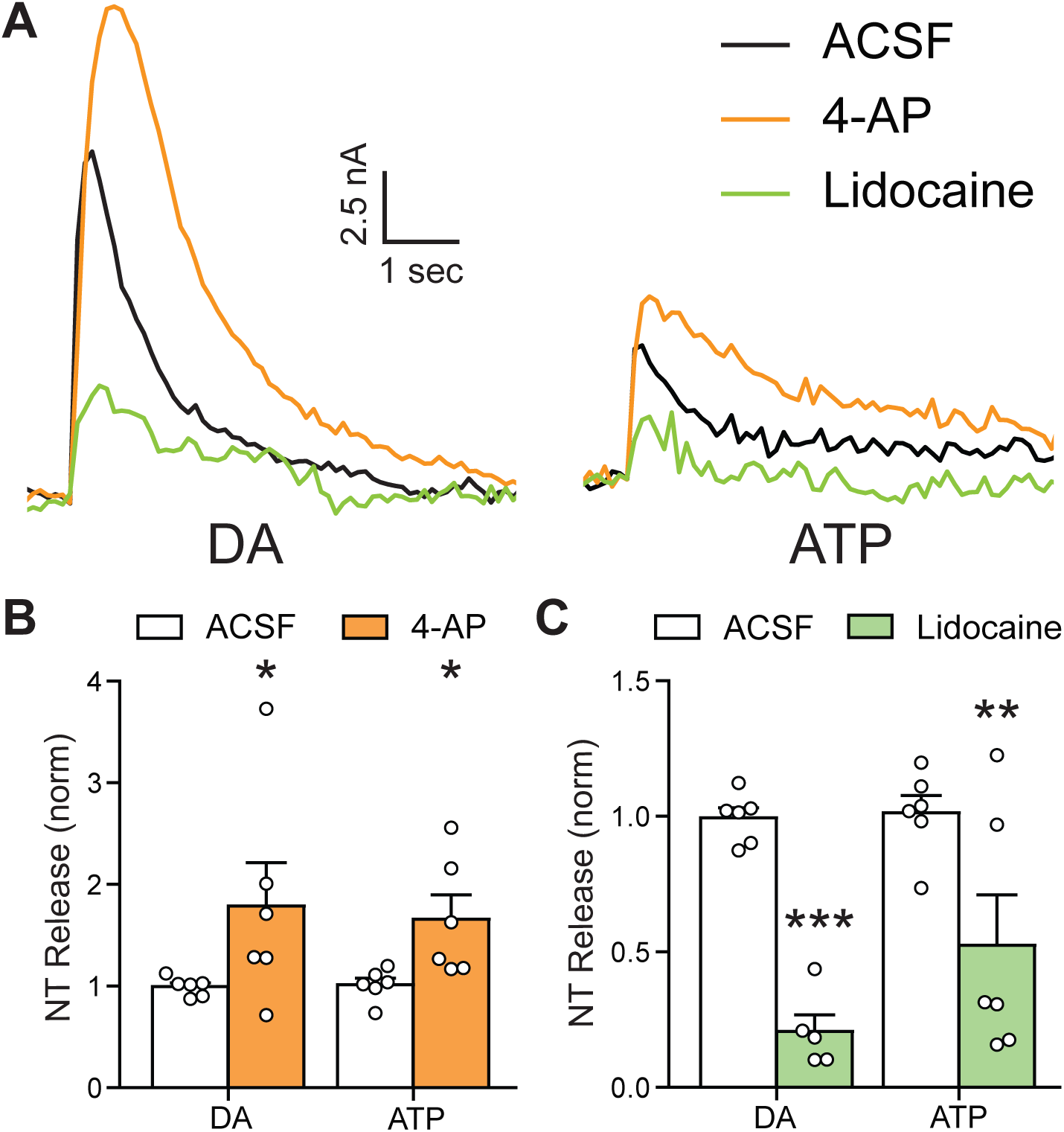
The Role of Action Potentials on Transmitter Release. **A** Representative traces of evoked DA and ATP release during ACSF (black), bath application of 4AP (30 μM, orange), and bath application of lidocaine (100 μM, green). **B** 4AP significantly increased the amplitude of evoked DA and ATP release. **C** Lidocaine significantly decreased the amplitude of evoked DA and ATP release compared to ACSF. Asterisks ** and *** indicate significance levels *p*<0.01 and *p*<0.001, respectively.

### D2 Receptor Activation Inhibits DA and ATP Release

Dopamine neurons express DA D2 autoreceptors, which regulate DA release through activation of inhibitory currents and reduced vesicular fusion (Ford, 2014). A submaximal concentration of the D2 receptor agonist quinpirole (100 nM) was bath-applied for one hour followed by application of the antagonist sulpiride (600 nM) (**Fig. 3**). Representative current tracings for evoked DA and ATP release are shown (**Fig. 3A**). Quinpirole administration significantly decreased electrically evoked DA and ATP release amplitude (**Fig. 3B**; two-way repeated-measures ANOVA; quinpirole: F_1,24_=45.94, *p*<0.0001, neurotransmitter: F_1,24_=0.6066, *p* =0.4437; interaction: F_1,24_=1.141, *p*=0.2961; Bonferroni post hoc, *p*<0.001 for DA and ATP). Sulpiride reversed this decrease, returning ATP and DA release amplitude to pre-quinpirole levels (**Fig. 3C**; two-way repeated-measures ANOVA; sulpiride: F_1,20_= 0.5713, *p*=0.4586; neurotransmitter: F_1,20_= 0.1165, *p*=0.7364; interaction: F_1,20_=0.1043, *p*=0.7501). These data show that ATP is regulated by inhibitory D2 receptors similarly to DA release.

**Figure 3:**
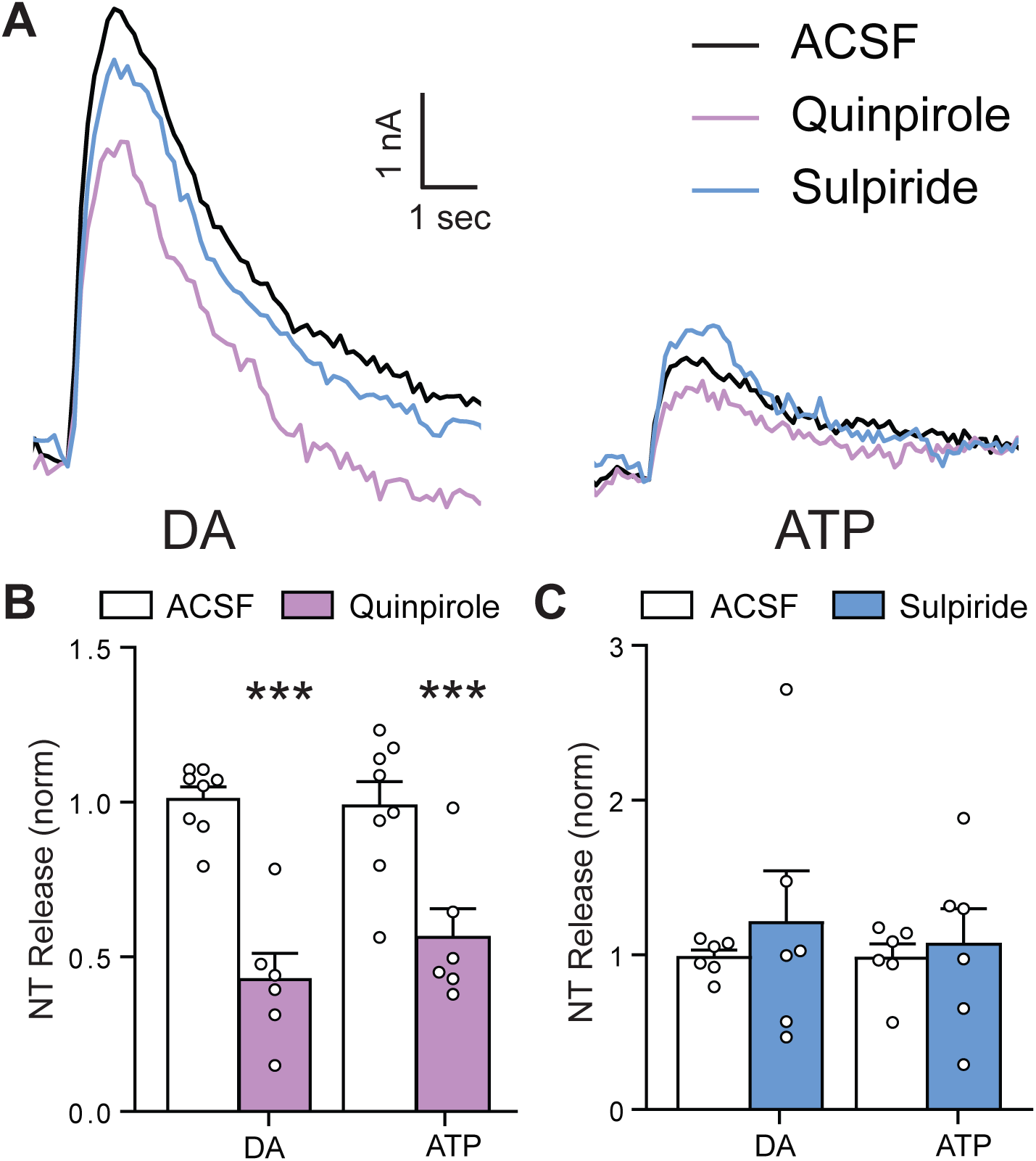
D_2_ Receptor Modulation and Effects on Release. **A** Representative traces of evoked DA and ATP release during ACSF (black), bath application of quinpirole (100 nM, purple), and bath application of sulpiride (600 nM, blue). **B** Quinpirole significantly decreased the amplitude of evoked DA and ATP release. **C** Sulpiride reverses the effects of quinpirole on DA and ATP release. Asterisks *** indicate significance level *p*<0.001 compared to ACSF.

### ATP Packaging is Dependent on DA Packaging

The vesicular monoamine transporter (VMAT) packages DA into synaptic vesicles (Krantz et al., 1997) and the VMAT blocker reserpine interrupts DA vesicular packaging, resulting in decreases in evoked DA release over a ∼1-hr period (Yorgason et al., 2017a). Reserpine (1 µM) was applied for 1 hour to test the relationship between DA and ATP vesicular packaging (**Fig. 4**). Importantly, vesicular depletion is dependent on activity/release, and stimulating release can facilitate more rapid DA depletion (Yorgason et al., 2017a). If the DA depletion is coupled to ATP depletion, then 4-AP should similarly decrease ATP. However, if ATP release is unaffected or enhanced by 4-AP, then this would suggest an uncoupling of vesicular packaging mechanisms. Thus, in these depletion experiments, 4-AP (30 µM) was consecutively added to the existing reserpine solution to accelerate DA depletion and test if ATP would be similarly depleted. Finally, lidocaine (100 µM) was added to the solution to block action potentials and assess VMAT-insensitive ATP stores. Example current tracings are shown for electrically evoked DA and ATP under the specific drug conditions (**Fig. 4A**). Reserpine administration significantly decreased DA release amplitude, which was further decreased by 4-AP or lidocaine administration (**Fig. 4B**; one-way repeated-measures ANOVA, F_3,9_=30.06, *p*<0.0001). Reserpine, 4-AP, and lidocaine administration each also significantly decreased electrically evoked ATP release amplitude (**Fig. 4C**; one-way repeated-measures ANOVA, F_3,9_=16.69, *p*=0.0005). Reserpine alone diminished ATP release by approximately 50%, demonstrating that ATP release is at least partially dependent on DA vesicle packing mechanisms. Importantly, 4-AP did not further decrease ATP release following reserpine, and unlike DA, ATP release continued to persist in the presence of lidocaine. This supports the existence of a conductive diffusion pathway of ATP release as previously described in other regions that is unaffected by blocking vesicular release machinery.

**Figure 4:**
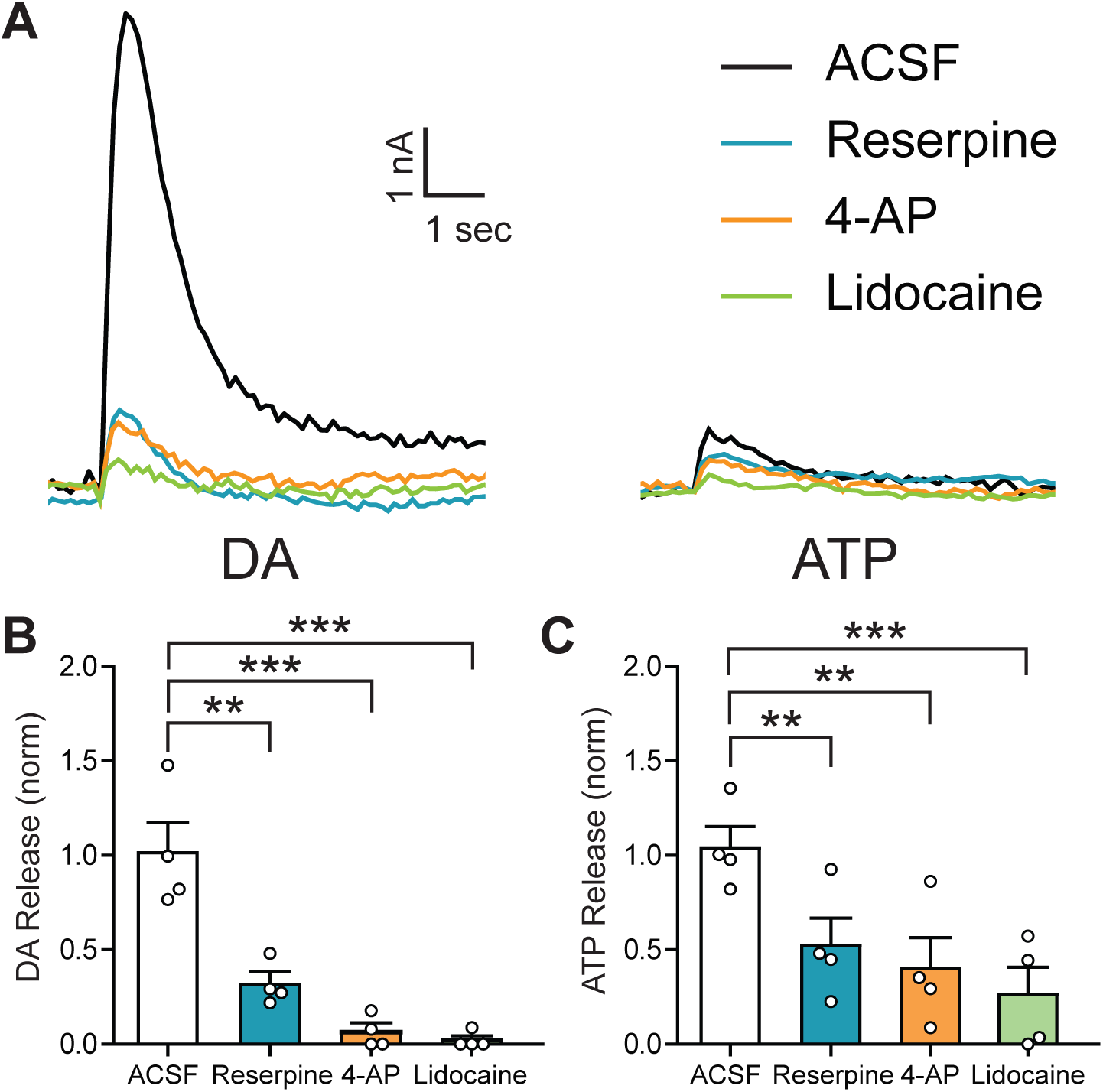
ATP and DA Vesicular Packaging: The Role of the VMAT. **A** Representative traces of evoked DA and ATP release during ACSF (black), bath application of reserpine (1 μM, cyan), 4AP (30 μM, orange), and lidocaine (100 μM, green). **B** Reserpine significantly decreases evoked DA release, which is then further reduced with 4AP and lidocaine (Tukey’s posttest). **C** Reserpine significantly reduced ATP release (Tukey’s post test) compared to ACSF, which was not further reduced by modulating action potentials. Asterisks *,*** indicate significance levels *p*<0.05 and *p*<0.001, respectively.

### ATP Release is Regulated by Nicotinic Acetylcholine Receptors

Cholinergic interneurons are powerful modulators of DA release (Yorgason et al., 2017a) and thus likely contribute to ATP release. Bath application of HEX (200 µM), a non-selective nAChR antagonist, was applied to NAc slices and DA and ATP release were measured (**Fig. 5**). Example current tracings are shown for electrically evoked DA and ATP before and after HEX (**Fig. 5A**). HEX reduced DA and ATP release amplitude (not shown; DA peak height: one-tailed paired t-test, *t*_9_=2.588, *p*=0.0147; ATP amplitude: two-tailed paired t-test, *t*_9_=2.821, *p*=0.02), and the change in amplitude was roughly similar between the two neurotransmitters (**Fig. 5B**; DA vs ATP amplitude change: two-tailed unpaired t-test, *t*_18_=1.408, *p*=0.1761). This change in release of DA and ATP after HEX application was significantly correlated between DA and ATP (**Fig. 5C**; Spearman r=0.6727, p=0.0390). The change in clearance (delta exponential decay, tau) of DA and ATP after HEX application was also significantly correlated (**Fig. 5D**; Spearman r=0.7333, p=0.0311). In line with our hypothesis of multiple ATP release mechanisms, blocking nAChR signaling reduced, but did not deplete, release of DA and ATP. This finding demonstrates that, like DA, vesicular ATP release is responsive to input from cholinergic interneurons and thus participates in mesolimbic circuitry implicated in disease states like addiction (Wadsworth et al., 2023; Yorgason et al., 2015, 2017).

**Figure 5:**
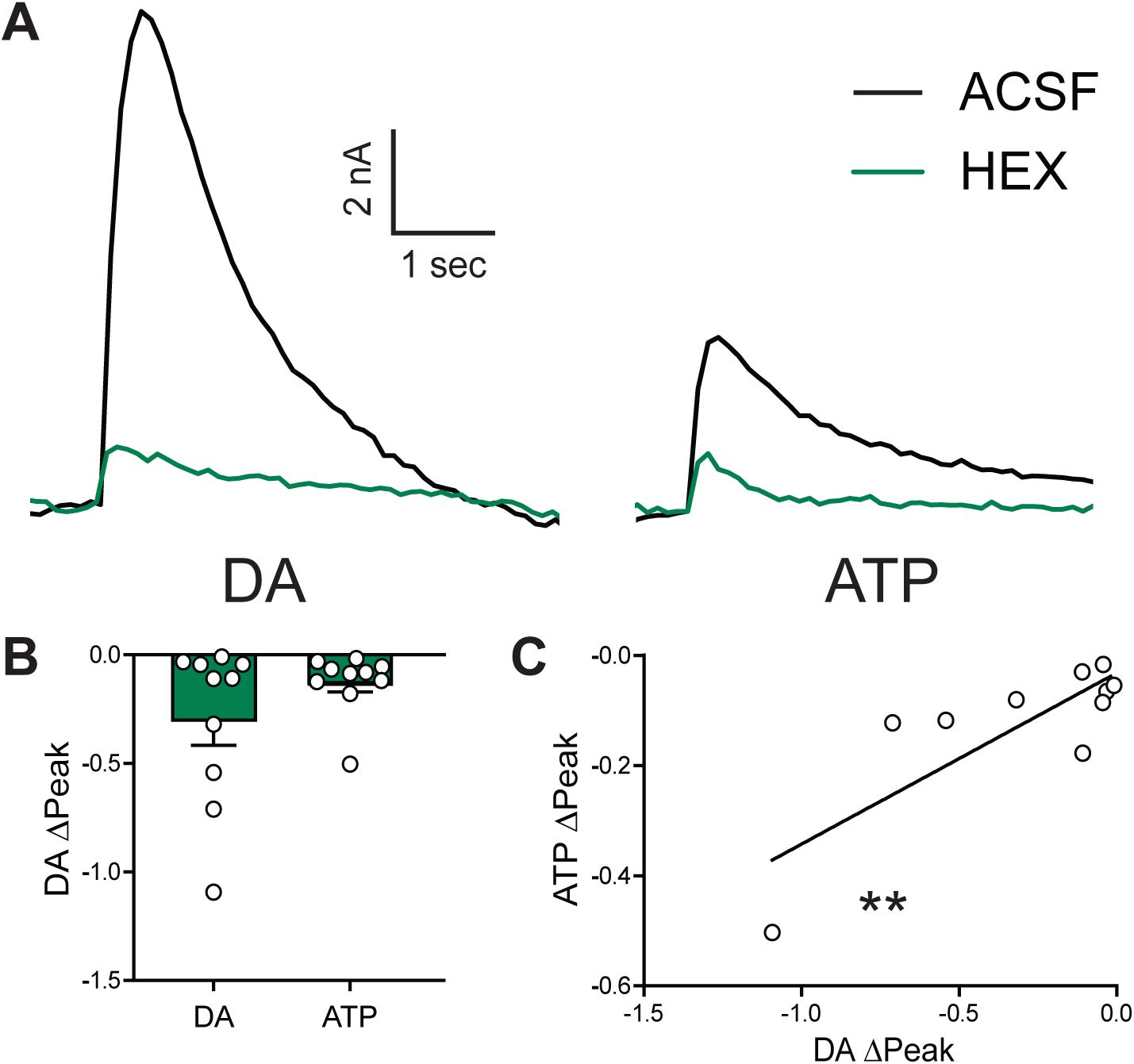
Cholinergic Modulation of ATP and DA Transmission via nAchR Inhibition. **A** Representative DA and ATP traces before (ACSF) and after hexamethonium (HEX; 200µM). HEX inhibits both evoked DA and ATP release. **B** Group data comparing HEX effects on DA and ATP release (*p*=0.1761). **C** Regression analysis establishing association between HEX effects on DA and ATP release (Spearman r=0.6727, p=0.0390), which indicates that HEX effects on DA are to a greater extent than that of ATP (i.e. the slope of the regression is different from 1) compared to ACSF. Asterisks *,**,*** indicate significance levels *p*<0.05, *p*<0.01, and *p*<0.001, respectively.

### Synaptic Reserve Vesicles Recruited by Cocaine Contain ATP

Cocaine is primarily known as a DA transporter blocker, but it also has effects on DA release. At low-to-middling doses, cocaine increases release of DA by mobilizing a reserve pool of DA-containing vesicles (Venton et al., 2006). However, at high doses, cocaine has been shown to decrease DA release due to off-target blockade of voltage-gated sodium channels (lending to its use as a local analgesic; Steffensen et al., 2008). We applied a high concentration of cocaine (30 µM) but were able to compare the effects of DAT blockade on DA and ATP transmission under both release conditions by analyzing data from different timepoints in the cocaine administration. Analysis of earlier timepoints shown here represents DA release-increasing effects of cocaine, where the concentration of cocaine was still low yet steadily increasing in the bath (**Fig. 6**). Shown are representative examples from two experiments (**Fig. 6A**) before and immediately after cocaine application. Both DA and ATP release transiently increased after cocaine superfusion (**Fig. 6B**; DA amplitude: one-tailed unpaired t-test, *t*_10_=2.299, *p*=0.0222; ATP amplitude: one-tailed unpaired t-test, *t*_10_=0.2845, *p*=0.0087; DA vs ATP amplitude change: *t*_5_=4.266, *p*=0.008). Cocaine superfusion at early timepoints amplified ATP release by 217±37% and DA release by 256±50%. Though the difference in ATP and DA release amplification did not reach statistical significance (not shown; one-tailed paired t-test, t_5_=1.614, p=0.0837), cocaine effects on ATP and DA release were strongly correlated (**Fig. 6D**; Pearson r=0.8876, *p*=0.0182, linear regression; F_1,4_=14.86). Clearance of both DA and ATP were reduced (**Fig. 6C**; DA tau: one-tailed unpaired t-test, *t*_10_=3.438, *p*=0.0032; ATP tau: two-tailed unpaired t-test, *t*_10_=4.485, *p*=0.0012). In contrast to release measures, cocaine effects on clearance immediately following its application were not significantly correlated between the two neurotransmitters (**Fig. 6E**; Pearson r=0.2347, *p*=0.6544, linear regression; F_1,4_=0.2332).

**Figure 6:**
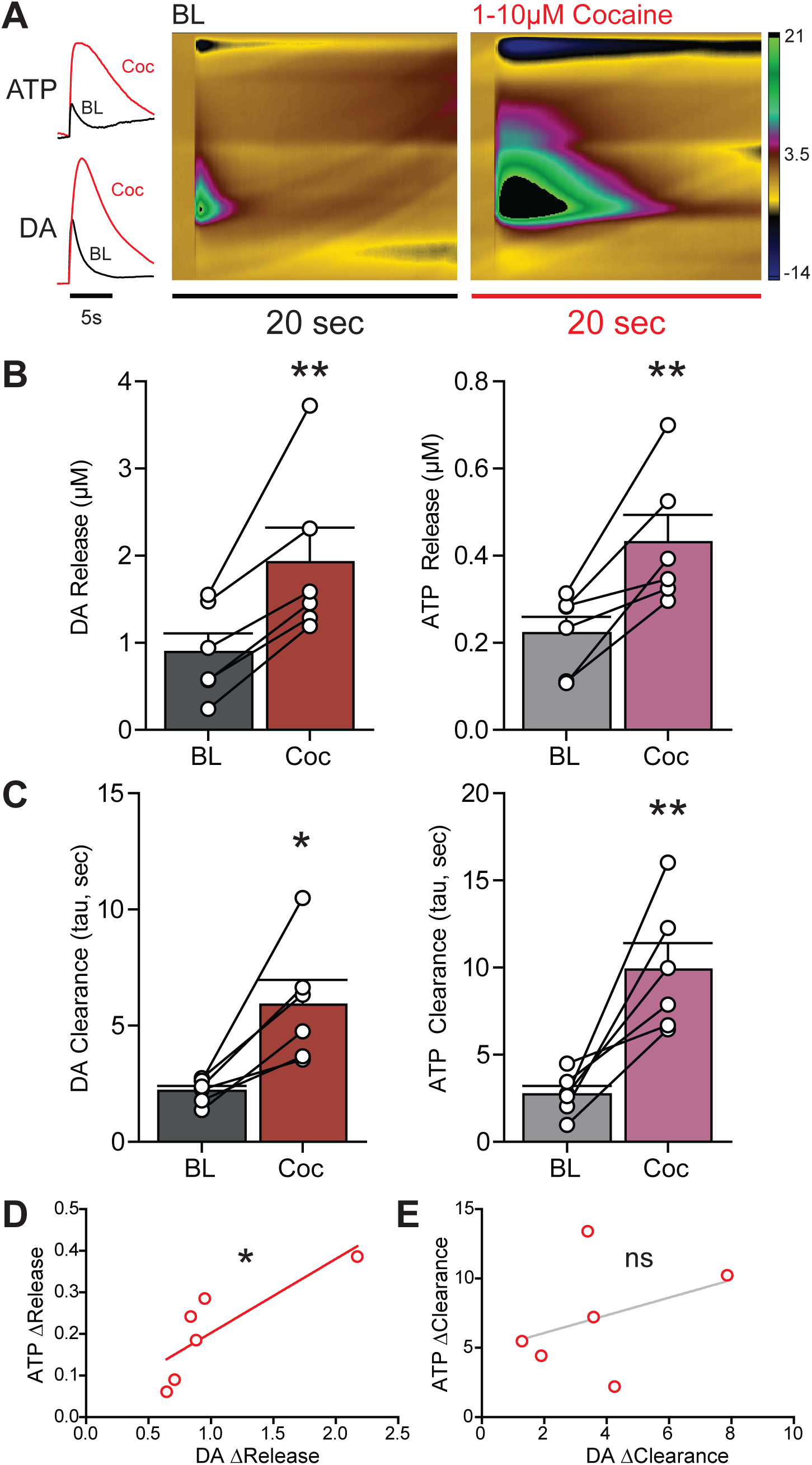
Synaptic Reserve Vesicles Recruited by Cocaine Contain ATP. **A** Example traces and colorplot during cocaine onset (approx. 1-10 µM; Coc) from experiments testing effects of cocaine on DA and ATP transmission. **B** Group data comparing baseline (BL) and Coc induced increases in DA and ATP release and **C** clearance. **D** There was a strong relationship between the coc-induced increases in ATP release and Coc-induced DA release within the same brain slice. **E** There was no apparent relationship between Coc-induced reductions in clearance between transmitters. Asterisks * and ** indicate significance levels *p*<0.05 and *p*<0.01, respectively.

### ATP Clearance is Attenuated by DAT Inhibition

The analysis of DA and ATP transmission described above was repeated for later timepoints where the concentration of cocaine was at its highest (30 µM) and the DA release-decreasing effects of high-dose cocaine were observed (**Fig. 7**). Shown are representative examples from two experiments (**Fig. 7A**) before and after cocaine was applied with sufficient time to reach the full dose. At 30 µM, the cocaine-induced increase in release seen previously (Fig. 6B) was abolished for both DA and ATP (**Fig. 7B**; DA amplitude: one-tailed paired t-test, *t*_5_=1.285, *p*=0.1275; ATP amplitude: two-tailed paired t-test, *t*_5_=0.2736, *p*=0.7953; DA vs ATP normalized amplitude change (not pictured): two-tailed paired t-test, *t*_5_=2.538, *p*=0.0520). Clearance of both neurotransmitters, however, remained impaired at this high concentration (**Fig. 7C**; DA tau: one-tailed paired t-test, t_5_=2.765, *p*=0.0198; ATP tau: two-tailed paired t-test, t_5_=4.088, *p*=0.0095). 30 µM cocaine effects on ATP and DA release were correlated (**Fig. 7D**; Pearson r=0.873, *p*=0.023). In contrast to Fig. 6E, clearance of ATP and DA at this dose was also correlated (**Fig. 7E**; Pearson r=0.913, *p*=0.011).

**Figure 7:**
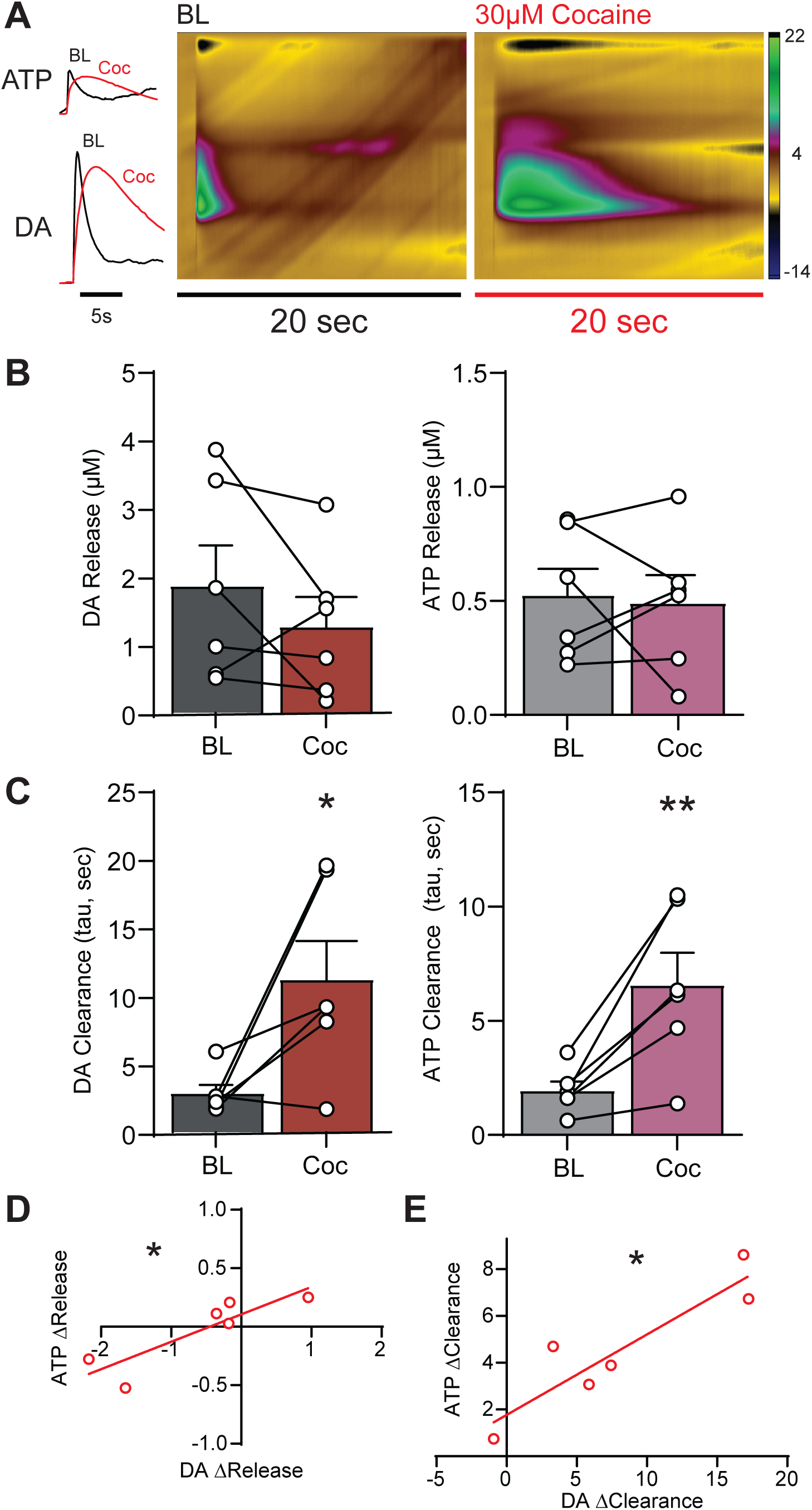
ATP Clearance is Attenuated by DAT Inhibition. **A** Example traces and colorplot at full cocaine concentration (30 µM; Coc) from experiments testing effects of cocaine on DA and ATP transmission. **B** Group data comparing baseline (BL) and Coc induced increases in DA and ATP release and **C** clearance. **D** There was a strong relationship between the Coc-induced increases in ATP release and Coc-induced DA release within the same brain slice. **E** Coc-induced reductions in clearance correlated between transmitters. Asterisks * and ** indicate significance levels *p*<0.05 and *p*<0.01, respectively.

## Discussion

### NAc ATP Release Mechanisms

All synaptic vesicles contain some ATP required for protein function, but the concentration of ATP within them can vary widely, and some have even been observed to contain only ATP (Pankratov et al., 2006). This and rapid synaptic clearance are characteristic of classical neurotransmitters like DA, and our observation that DA and ATP have strongly correlated reuptake rates indicates that ATP has neurotransmitter-like function (**Fig. 1**). This supports the growing evidence that ATP co-releases with DA through calcium-dependent vesicular exocytosis (Borgus et al., 2021; Ho et al., 2015; Nguyen et al., 2015). In the striatum, most secretory vesicles contain a high concentration of ATP, suggesting that a major mechanism for releasing ATP as a neurotransmitter in this region is through vesicular co-release with other signaling molecules, including DA (Borgus et al., 2021; Estévez-Herrera et al., 2016). Although ATP vesicular packaging is fairly ubiquitous, it is also noteworthy that the expression of VNUT affects ATP vesicular packaging (Estévez-Herrera et al., 2016), indicating that there are likely neuronal subtype differences in vesicular ATP packaging and release.

Blocking Kv channels on the pre-synaptic membrane with 4-AP caused more DA and ATP to be released, an effect that was reversed when NaV channels were blocked by lidocaine (**Fig. 2**). This suggests that both DA and ATP evoked release are dependent on an action-potential mediated mechanism. Blocking potassium-driven membrane repolarization also permits an increased amount of calcium to enter through voltage-gated calcium channels, activating synaptotagmin and other calcium-dependent vesicular release proteins (Körber & Kuner, 2016; Yang et al., 2020). ATP and DA are released in greater amounts when these channels are blocked, so they likely rely on a vesicular release mechanism when electrically evoked. Similarly, decreased levels of DA and ATP release following lidocaine administration could be attributed to action potential-mediated effects on release, supporting a vesicular release mechanism for evoked ATP release. These experiments show that DA and ATP release are mediated through similar action potential-dependent mechanisms. Of note, we observed that ATP exhibited a slower release velocity and that some ATP release was insensitive to lidocaine (therefore independent of action potentials). These results show that there is likely a subset of ATP release occurring through conductive mechanisms, such as pore formation connexins and pannexin-1 (Cisneros-Mejorado et al., 2015; Xia et al., 2012).

### D2 Receptor Activation Inhibits DA and ATP Release

Activating D2 receptors with quinpirole significantly decreased evoked ATP and DA release amplitude, an effect that was reversed by administering a D2 receptor antagonist sulpiride (**Fig. 3**). This finding is consistent with the current understanding of DA autoregulation, which has been thoroughly characterized (Heidbreder & Baumann, 2001; C. Liu & Kaeser, 2019; Sulzer et al., 2016), and additionally suggests that D2 autoreceptors regulate evoked release of both DA and ATP. The D2 autoreceptor is known for mediating critical inhibitory feedback, whereby impaired D2 receptor function leads to abnormally elevated dopamine levels observed in several neuropsychiatric disorders. For instance, loss of D2 receptor function has been observed in DAT-KO mice, and the subsequent prolonged elevations in DA during critical developmental periods are associated with increased anxiety-like behaviors and vulnerability to psychostimulant abuse (Jones et al., 1999; Yorgason et al., 2016). Our observation that D2 receptors acutely regulate extracellular ATP alongside DA warrants future investigations into whether D2 receptor modulation of ATP transmission plays an undiscovered role in these pathological states.

Based on prior studies of DA release inhibition, the downstream effects of D2 activation most likely responsible for ATP inhibition are direct mechanisms in membrane hyperpolarization, including activation of voltage gated K+ channels (Fulton et al., 2011; Neve et al., 2004). G-protein coupled inward rectifying K+ (GIRK) channels have also been implicated in DA somatic activity, but do not appear to play a substantial role in D2 autoinhibition at NAc terminals (Martel et al., 2011). However, considering the importance of D2-GIRK activity in the soma, future studies might expect to find a major role for GIRKs in ATP release in midbrain regions projecting to the NAc. As D2 receptor dysfunction is also thought to underlie changes in DA synthesis and packaging mechanisms (Jones et al., 1998, 1999), another avenue of investigation could determine whether D2 receptor activation induces lasting changes in terminal VNUT expression and associated ATP release, particularly in early life stress or substance use models.

### ATP Release is Dependent on DA Packaging

DA and ATP molecules have previously been shown to form complexes in vitro (Berneis et al., 1971; Granot & Fiat, 1977; Taleat et al., 2018a; Weder & Wiegand, 1973). Though this has not yet been demonstrated in vivo, ATP has been observed in high concentrations in synaptic vesicles, and the inner vesicular pH is theoretically well-suited to ATP-DA complex formation (Estévez-Herrera et al., 2016). If these complexes do form in brain tissue, they could serve to increase vesicular packaging of these transmitters by facilitating higher packing density. Indeed, there is already evidence that ATP enhances vesicular packing efficiency and release quanta of catecholamines (Estévez-Herrera et al., 2016; Larsson et al., 2019; Majdi et al., 2019).

We tested whether NAc ATP release is dependent on successful packaging of DA into vesicles and found that blockade of VMAT (which packages DA) with reserpine reduced both DA and ATP release (**Fig. 4**). The incomplete reduction in DA release was expected since total depletion of DA release via reserpine is stimulation-dependent (Jones et al., 1998; Karkhanis et al., 2019; Yorgason et al., 2015). However, the observation that ATP release was VMAT-dependent is notable as it adds novel context to previous studies where knockdown or knockout of VNUT (which packages ATP into synaptic vesicles) likewise resulted in decreased quantal release of both ATP and catecholamines (Borgus et al., 2021; Estévez-Herrera et al., 2016; Miras-Portugal et al., 2019; Sakamoto et al., 2014). Taken together with previous work, our result demonstrates that vesicular concentrations of ATP and DA are mutually dependent. While we and others suspect that this improvement in packing efficiency owes to the formation of “colligative” DA-ATP complexes in vesicles (Estévez-Herrera et al., 2016; Taleat et al., 2018), additional studies will be necessary to conclusively demonstrate the existence of such complexes in vivo. A definitive approach might be to directly image these complexes, but this would likely require an expensive high-resolution imaging technique like cryo-electron microscopy—the chemical associations attributed to the formation of these small-molecule complexes are weak and could potentially be interrupted by fluorophore binding. Nonetheless, fluorescence microscopy has been used to visualize ATP in subcellular structures, and new fluorescent probes for ATP continue to be developed (Corriden et al., 2007; Imamura et al., 2009; Sørensen & Novak, 2001; Tan et al., 2017). Of interest, recent advances in electrochemistry have succeeded in creating carbon nanoelectrodes capable of analyzing the content of single vesicles through fast scan cyclic voltammetry, intracellular vesicle impact electrochemical cytometry, and other methods (Li et al., 2015; Roberts et al., 2020; for review, see Y. Liu et al., 2021). These techniques enable simultaneous quantification of DA and ATP molecules within a single vesicle, which could be used to assess packing efficiency. They might also prove useful in confirming the relative contribution of vesicular ATP release to the total amount of extracellular ATP release.

Following reserpine administration, both 4-AP and lidocaine conditions significantly decreased evoked DA release and nearly abolished the signal entirely. In contrast, 4-AP and lidocaine decreased ATP release in some but not all experiments, providing yet further evidence of both action potential dependent and independent ATP release. These results are consistent with the reported role of conductive ATP release in establishing “basal” cell signaling (Hussl et al., 2007; Ostrom et al., 2000; for review, see Corriden & Insel, 2010). However, non-exocytotic ATP release also occurs in response to mechanical stress or cellular swelling; this has been well-described in neuronal and non-neuronal tissue alike (Enomoto et al., 1994; Hazama et al., 2000; Lee et al., 2022; Mitchell et al., 1998; Osipchuk & Cahalan, 1992; Sprague et al., 1998; Xia et al., 2012). An important caveat of our technique is that the recording and stimulating electrodes are inserted directly into the brain slice, piercing the tissue and causing a small amount of damage. Thus, we speculate that some amount of the conductive ATP release we observed was cell signaling triggered by mechanical stress. Applying light- or chemical-based techniques without invasive mechanical manipulations in future studies will help differentiate between these two sources of conductive ATP release and verify their functional roles in NAc terminals.

ATP is a known Damage-Associated Molecular Pattern (DAMP), and extracellular ATP induces local immune responses such as activation of microglia to release cytokines and repair tissue (Cauwels et al., 2014; Davalos et al., 2005; Murana et al., 2017). We and others have previously observed robust regulation of DA release by cytokines (Kutlu et al., 2018; Payne et al., 2026; Ronström et al., 2023), implicating these neuroimmune interactions as a mechanism through which extracellular ATP can indirectly affect DA release. Additionally, ATP release can indicate and repress excessive excitatory activity by acting as an autocrine signal through activation of K_ATP_ channels (de Siqueira et al., 2025) or through conversion to adenosine and resultant effects on inhibitory GPRCs (Borgus et al., 2021; Lazarus et al., 2011; Lee et al., 2022; Pajski & Venton, 2010; Rosin et al., 1998). The former of these two is of particular importance since DA terminals are already known to express autoregulatory K_ATP_ channels (Lee et al., 2022; Martel et al., 2011; Patel et al., 2011). Besides DA terminals, K_ATP_ channels are also expressed on local ATP receptors, including purinergic receptors on local microglia (Kitaoka, 2023; Wadsworth et al., 2024), A2A receptors on medium spiny neurons, and A1A receptors on cholinergic interneurons (Lazarus et al., 2011; Rosin et al., 1998). We have previously shown considerable regulation of DA release by local cholinergic interneurons (Brundage et al., 2022; Gao et al., 2019; Wadsworth et al., 2023, 2024; Yorgason et al., 2017, 2022), so ATP transmission might also affect DA release through a cholinergic mechanism in addition to the mechanisms described above.

### ATP Release is Regulated by Cholinergic Receptors

Activation of nAChRs, which are commonly expressed on presynaptic DA terminals in the NAc, induces DA release (Koranda et al., 2014; Yorgason et al., 2017). Application of HEX, a nAChR blocker, reduced both DA and ATP release, demonstrating that nAChR-expressing DA terminals participate in ATP-DA corelease (**Fig. 5**). Interestingly, previous work in different neuronal cell types has found that nAChRs and P2X receptors colocalize on the plasma membrane and cross inhibit one another (Decker & Galligan, 2009; Khakh et al., 2005). If this phenomenon is found in NAc terminals as well, it would lend further support to our hypothesis that cholinergic interneurons are one of the inhibitory feedback mechanisms through which ATP release from DA terminals downregulates neurotransmitter release. We also observed that ATP release was less influenced by nAChR blockade than DA. Modulation of DA release by nAChRs is action potential-dependent (Yorgason et al., 2017), so this result is consistent with our previous observations of action potential-independent ATP release. It is also likely that ATP transmission occurs in other terminals expressing few or no nAChRs, possibly including non-dopaminergic terminals, which should be explored in future studies.

### Synaptic Reserve Vesicles Recruited by Cocaine Contain ATP

Bath application of cocaine affected release and clearance of DA and ATP similarly. Specifically, cocaine induced an acute increase in both DA and ATP release (**Fig. 6**) that subsided once the full concentration was reached (**Fig. 7**). This release effect was well correlated at both early and late timepoints. We considered two possibilities most likely to cause this result. One explanation is that reserve vesicles recruited by cocaine contain ATP in addition to DA, while the other is that ATP release through conductive mechanisms increases in response to downstream effects of cocaine. Both explanations would be in line with the described role of ATP as a cellular stress signal: Cocaine increases DA release by mobilizing synapsin-dependent intracellular vesicular reserve pools (Venton et al., 2006), and several studies have shown neuroprotective effects of ATP release in response to excessive firing (de Siqueira et al., 2025; Fujikawa et al., 2012; Xiong et al., 2022; Yamazaki & Fujii, 2015). However, we reasoned that a higher concentration of cocaine would be expected to further increase reserve vesicle release demand, cellular stress, and subsequent damage signaling by ATP. Since ATP release amplitude decayed alongside that of DA when cocaine concentration was high, the first explanation (that ATP is released from reserve vesicles) is more likely. This conclusion is further reinforced by the fact that cocaine blocks voltage-gated sodium channels at high doses (Steffensen et al., 2008), which would interrupt action potential-dependent vesicular release but not conductive diffusion pathways. Thus, the attenuated ATP release (relative to drug onset) at high cocaine concentration again indicates the presence of exocytotic ATP release and additionally implicates synapsin as a potential target for elucidating this mechanism.

### ATP Clearance is Attenuated by DAT Inhibition

Although the change in clearance rates between neurotransmitters only correlated once the high dose of cocaine was achieved, cocaine slowed clearance of both neurotransmitters at both analyzed time points. This result introduces unexpected evidence for ATP clearance through DAT-related mechanisms. Since ATP and DA form complexes in solution (Taleat et al., 2018), one intriguing possibility is that individual complexes of DA and ATP molecules are absorbed by DATs as a single unit, so inhibiting DAT blocks the joint reuptake of both molecules. Paired with the observed correlations in release, this could also provide a mechanism for co-packaging of ATP into vesicles alongside DA as a single complex. However, whether such complexes form in biological tissue has not yet been tested, and the chemical strength of the association between ATP and DA molecules is reportedly weak, so this explanation is less persuasive. Despite both molecules exhibiting sensitivity to DAT blockade, the clearance of the two transmitters does not appear to be strongly related since correlation was only observed at the high concentration. Downstream consequences of DAT inhibition are therefore more likely to mediate this relationship. This is not too surprising since there are several other mechanisms available for ATP and DA clearance alike. The key ATP clearance proteins that have been identified include adenosine kinase, equilibrative nucleoside transport 1, and adenosine deaminase (Nguyen et al., 2015; Yorgason et al., 2017). Whether these proteins contribute to NAc ATP synaptic clearance remains unknown, so additional experiments are needed to understand the link between DA and ATP clearance.

## Conclusion

To our knowledge, we are the first to demonstrate ATP release from DA terminals in the nucleus accumbens and to characterize its mechanism. This work provides several lines of converging evidence for action potential-dependent vesicular ATP co-release alongside DA in the NAc. ATP release and clearance were tightly coupled to that of DA, with ATP releasing in smaller concentrations than DA. Cocaine, a DAT inhibitor that is also known to increase vesicular DA release, increased release of both DA and ATP. This increase was strongly correlated between the neurotransmitters. Further, both DA and ATP release were affected by manipulation of D2Rs with quinpirole and sulpiride, and 4-AP and lidocaine experiments demonstrated that release of both neurotransmitters is action-potential dependent. Notably, while DA levels became undetectable under lidocaine conditions, ATP release persisted (albeit in a smaller quantity), suggesting that some ATP is also released through non-exocytotic mechanisms. Manipulations of vesicular packaging and NaV channel function lend further support to this idea. Both reserpine and lidocaine prevented the 4-AP-induced increase in ATP to instead decrease (without fully preventing) ATP release, and application of cocaine at a concentration high enough to block voltage-gated sodium channels similarly reduced but did not abolish the ATP signal. This is also supported by our investigation of cholinergic regulation of ATP release. HEX blocked both DA and ATP release, but ATP release was less affected than DA. Our experiments indicate that ATP is packaged into vesicles (including synaptic reserve vesicles) alongside DA and exocytotically released, with an undercurrent of conductive ATP release that is not dependent on action potentials. Together, this demonstrates that ATP in the NAc acts through both vesicular and non-vesicular mechanisms to acutely regulate DA transmission.

## Acknowledgements

We would like to acknowledge internal funding from Brigham Young University andLPHS NIH grants AA030577 to JTY, and AA020919 and DA035958 to SCS. The authors have no competing conflicts of interest to disclose.L

